# Proteomic Characterization of the Venom Of Five *Bombus (Thoracobombus)* Species

**DOI:** 10.1101/193524

**Authors:** Nezahat Pinar Barkan, Mustafa Bilal Bayazit, Duygu Demiralp Özel

**Affiliations:** Hacettepe University, Faculty of Science, Department of Biology, Ankara, Turkey; University of Lausanne, Department of Fundamental Neurosciences, Lausanne, Switzerland; Ankara University, Faculty of Engineering, Biomedical Engineering, Ankara, Turkey

**Keywords:** Bumble bees, MALDI-TOF MS, Proteomics, 2D-PAGE, Venom

## Abstract

Venomous animals use venom; a complex biofluid composed of unique mixtures of proteins and peptides, to act on vital systems of the prey or predator. In bees, venom is solely used for defense against predators. However, the venom composition of bumble bees (*Bombus sp.*) is largely unknown. *Thoracobombus* subgenus of *Bombus sp.* is a diverse subgenus represented by 14 members across Turkey. In this study, we sought out to proteomically characterize the venom of five *Thoracobombus* species by using bottom-up proteomic techniques. We have obtained two-dimensional polyacrylamide gel (2D-PAGE) images of each venom sample. We have subsequently identified the protein spots by using matrix assisted laser desorption ionization / time of flight mass spectrometry (MALDI-TOF MS). We have identified 47 proteins for *Bombus humilis*; 32 for *B. pascuorum*, 60 for *B. ruderarius*; 39 for *B. sylvarum* and 35 for *B. zonatus*. Our analyses provide the primary proteomic characterization of five bumble bee species’ venom composition.

## Introduction

In venomous animals, the use of venom or poison is one of the most essential mechanisms used for either capturing prey or defending oneself against predators. In competing for resources, venom is an adaptive trait and an example of convergent evolution. It is a complex biofluid secreted by venom gland and is composed of unique mixture of proteins, enzymes, peptides and biogenic amines that act on vital systems of the recipient (Calvete, 2009). In real bees (Apoidea), largest family of bees with over 20.000 members (Fitzgerald, 2013), venom is solely used for defense against predators (Arbuckle, 2017).

Among Apoidea, the majority of venom studies are focused on honey bee *Apis mellifera* (*A. mellifera). Bombus sp.,* which also belong in the same family as *A. mellifera,* are pollinators for various native and cultivated plants (Free, 1993). The venom composition of *Bombus sp.* is poorly understood.

Previous studies on *Bombus sp.* venom have mainly focused on characterization and isolation of highly abundant single molecules using bio-assays and chemical sequencing via Edman degradation (Choo, Lee, Yoon, Je, et al., 2010; Choo, Lee, Yoon, Kim, et al., 2010; Choo, Lee, et al., 2012; Choo et al., 2011; Choo, Yoon, & Jin, 2012; Favreau et al., 2006; Kim et al., 2013; Qiu et al., 2011; Qiu, Choo, Yoon, & Jin, 2012a, 2012b; Qiu et al., 2013; Wan et al., 2014). Among the described peptides, bombolitins are unique to *Bombus sp.* and enhance phospholipase A2 in liposome hydrolysis, in a similar fashion to melittin found in *A. mellifera* venom and crabrolin and mastoparan found in vespid *Vespa crabro* venom (Argiolas & Pisano, 1985). Phospholipase A2 (Hoffman, El-Choufani, Smith, & De Groot, 2001; Hoffman & Jacobson, 1996; Van Vaerenbergh, Debyser, Smagghe, Devreese, & de Graaf, 2015), serine proteases (Choo, Lee, Yoon, Kim, et al., 2010; Choo et al., 2011; Kim et al., 2013; Qiu et al., 2011; Qiu et al., 2012b; Van Vaerenbergh et al., 2015), serine protease inhibitors (Qiu et al., 2013; Wan et al., 2014), acid phosphatases (Hoffman et al., 2001; Hoffman & Jacobson, 1996; Van Vaerenbergh et al., 2015), arginine kinase (N. P. Barkan, Demiralp, & Aytekin, 2015), hyaluronidase (Hoffman & Jacobson, 1996), putrescine (Tom Piek, 2013), citrate (Fenton et al., 1995), defensin (Rees, Moniatte, & Bulet, 1997), and acethylcoline (T Piek, Veldsema-Currie, Spanjer, & Mantel, 1983) have also been isolated from *Bombus sp*. venom.

Recently, the whole genome sequencing of European large earth bumblebee *B. (Bombus) terrestris* has become publically accessible. The accessibility to the genome sequence, combined with the use of mass spectrometry, paved the way for deeply analyzing the venom proteome composition of this species. This study has identified 55 new molecules in the *B. terrestris* venom. Moreover, 72% of identified molecules had homologs in *A. mellifera* venom indicating that while there are some species-specific differences, these two bee species share major defense mechanisms. *B. terrestris* is the only member of the *Bombus sp.* whose venom proteome composition has been characterized.

In bumble bees, color patterns show similarity among species or are highly variable within species. Therefore, classification based on coat color can be unreliable (P. Williams, 2007). This is especially true for the *Thoracobombus* subgenus which consists of 14 species in Turkey that are morphologically difficult to separate. To overcome this, we have previously used landmark based geometric morphometrics on their wing structure (N. P. A. Barkan, A. M., 2013). Other approaches, use of pheromone analysis (Hovorka, Valterova, Rasmont, & Terzo, 2006; Terzo, Valterová, & Rasmont, 2007; Terzo, Valterova, Urbanova, & Rasmont, 2003; Urbanová, Valterová, Rasmont, & Terzo, 2002) and DNA based molecular techniques (S. Cameron, Hines, & Williams, 2007; S. A. Cameron, Hines, & Williams, 2006; S. A. Cameron & Williams, 2003; Du et al., 2015; Kawakita et al., 2003; Kawakita et al., 2004; P. H. Williams et al., 2012) have also been used in differentiating bumble bee species.

Turkey is regarded as one of the countries with the highest amount of species richness both in number and diversity in the West-Palaearctic region. The main aim of this study is to determine and compare the venom protein profiles of different *Thoracobombus* species of Turkey using bottom-up proteomic strategies. With the adoption of these approaches, not only will we characterize the venom proteome of *Thoracobombus* species for the first time, but we will also highlight the proteomic differences within the subgenus that may help in differentiating between the species.

## 2. Methods

### 2.1. Venom Extraction

*B. humilis* (Illiger), *B. pascuorum* (Scopoli), *B. ruderarius* (Müller), *B. sylvarum* (Linnaeus), and *B. zonatus* (Smith) were collected from Ankara, Bolu, Çankırı and Kayseri provinces from Middle Anatolian Region during June, July, August and September in 2013 and 2014 (Table S1), with the permission from the Republic of Turkey, Ministry of Forestry and Water Affairs Directorate of Nature Conservation & National Parks, and the Republic of Turkey, Ministry of Food, Agriculture & Livestock. Alive specimens were transported to Hacettepe University Department of Biology for venom extraction. For venom extraction, the venom reservoir is removed from the stinger in a sterile environment with forceps and the membrane is disrupted afterwards. From each individual, this method yielded 2-5 μl of venom which were collected in a cryotube and transported in liquid nitrogen. For proteomic analyses venom samples were pooled, while for spectroscopic analyses, populations were pooled separately. Collected samples were stored at - 80°C until use.

### 2.2. Proteomic Analysis

#### 2.2.1. Protein Quantification and Two-Dimensional Gel Electrophoresis

Venom of aforementioned bumble bee species was collected from 20 individuals per species. Protein concentrations of the venom samples were obtained by using the Bradford assay (Bradford, 1976). Venom was dissolved in 20 μl rehydration buffer [7M urea, 2 M thiourea, 4% CHAPS (w/v), 1% ampholytes (pH 3-10), 10 mM DTT and trace amount of bromophenol blue]. 2D-PAGE was performed as previously described (Igci & Demiralp, 2012) in three technical replicates and a total of 50 μg of protein was loaded per gel.

#### 2.2.2. 2D-PAGE Analysis

2D gel electrophoresis (2DE) images from venom samples of 5 different *Thoracobombus* species were analyzed using PDQuest Software Advanced 8.0.1 (Bio-Rad, CA, USA). PDQuest software examines each spot on every gel image and it picks a master gel containing most spots. All spots in each gel image is then matched onto the master gel. Each post is annotated with an identification number (SSP). For quantifying spots, four different normalization options are performed including total density in gel images, local regression, total quantity in valid spot and mean of log ratios. In the present study, 2DE image analysis is performed for both detecting common and/ or unique spots between samples and for quantifying spot intensity which accounts for the amount of protein in that spot. All spots were matched manually and ‘total density in gel images’ normalization was performed. Two-way ANOVA was performed to detect significancies in spot intensities (IBM Corp. Released 2010. IBM SPSS Statistics for Windows, Version 19.0. Armonk, NY: IBM Corp.).

#### 2.2.3. In-gel enzymatic digestion, Matrix-Assisted laser desorption ionization/ time of flight (MALDI-TOF) mass spectrometry and peptide mass fingerprinting (PMF) analysis

In gel trypsinization of protein spots was performed as previously described (Igci & Demiralp, 2012). Obtained peptides were loaded on MALDI-TOF plates using ZipTip^®^ Pipette Tips (Millipore). Tryptic peptides were dissolved in sample solvent containing 0.1% and 50% acetonitrile and mixed with an equal volume of matrix solution containing alpha-cyano-4-hydroxycinnamic acid, 0.1% TFA and 75% acetonitrile. 1.5 μl of these mixtures were spotted onto a target plate and analyzed with MALDI-TOF mass spectrometer (Water, UK) operated in positive-ion reflectron mode. Obtained spectra were then examined in MASCOT search engine (http://www.matrixscience.com/cgi/search_form.pl?FORMVER=2&SEARCH=PMF). Organisms taxonomically related to bumble bees were selected as bumble bees are not present in the database. “Other Metazoa” was selected as the taxonomical group as it is a diverse group including all venomous animals and *Drosophila* sp., the taxonomically closest species to bumble bees. Both Swissprot and NCBInr databases were used to maximize the number of hits. Identified proteins were categorized into protein families using Swissprot database.

## 3. Results

### 3.1. 2D-PAGE Analysis of *Thoracobombus* Venom

2DE gel images were obtained from each venom sample (Fig 1).

**Fig. 1.**
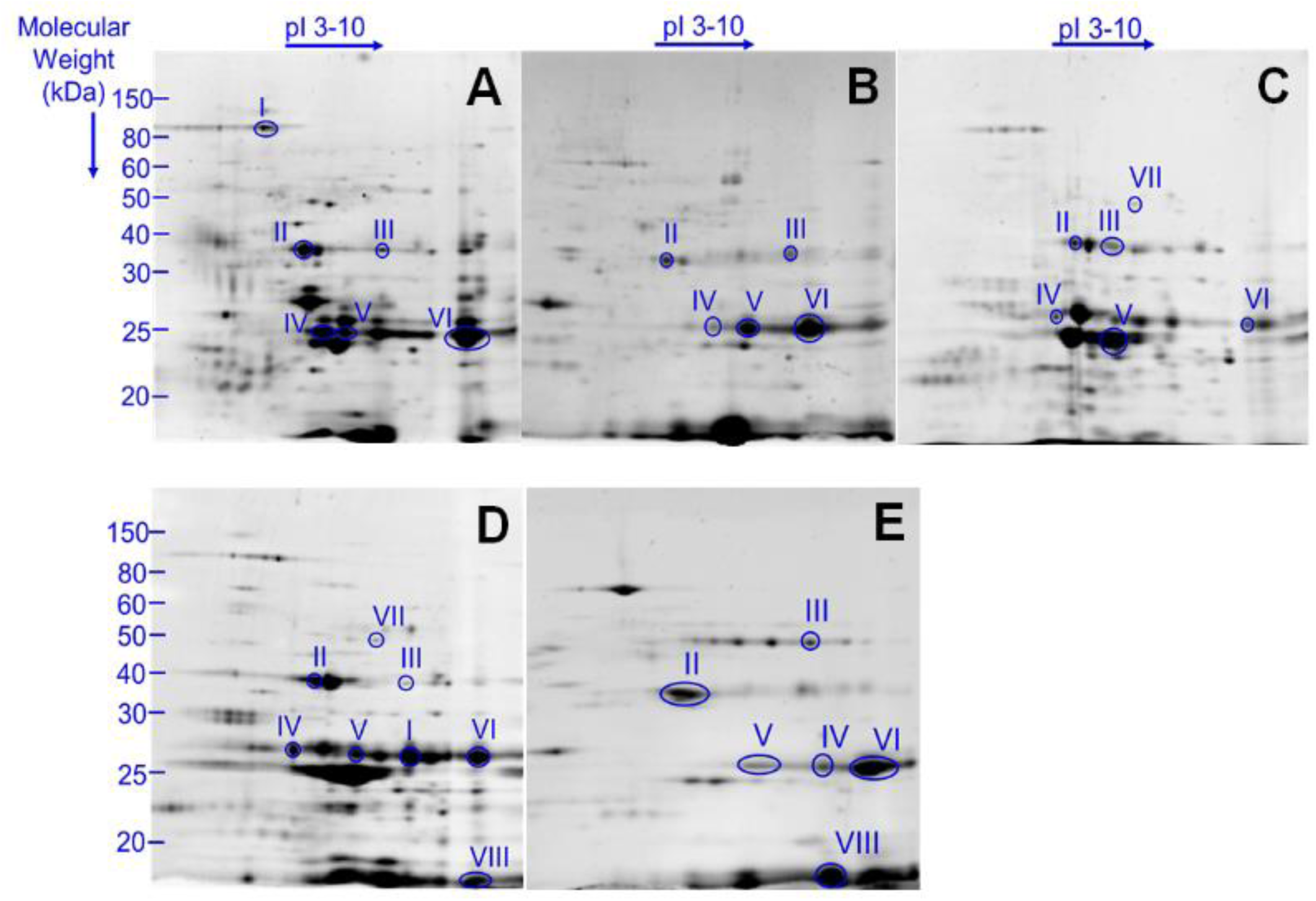
2DE gel images of *Thoracobombus* Venom. Representitive images of venom profile of (A) *B. humilis,* (B) *B. pascuorum* (C) *B. ruderarius* (D) *B. sylvarum* (E) *B. zonatus*. Gel analyses were conducted in triplicates. Putative venom toxins marked with roman numerals: I: Peptidase I*, Aminopeptidase N**, II: Venom acid phosphatase Acph1-like, III: Hyaluronidase 1, IV: Venom protease, V: Arginine Kinase, VI: Phospholipase A2, VII: Serine protease snake, VIII: Kunitz-type serine protease inhibitor. (See: Table 1).

**Table 1.** Putative toxins identified in *Thorocabombus* venom. Toxin names and the corresponding 2DE gel spots are indicated. The presence of the identified toxin (and/or its homologs/ortholog) in *B. terrestris and A. mellifera* venom is presented.

| Putative Toxin Family | Spot | Toxin Name | *B. terrestris* | *A. mellifera* |
| --- | --- | --- | --- | --- |
| *Peptidase/protease* | I | Peptidase I*,<br>Aminopeptidase N** | Dipeptidyl peptidase IV,<br>Serine carboxypeptidase,<br>Prolylcarboxypeptidase | Dipeptidyl peptidase IV,<br>Serine carboxypeptidase,<br>Prolylcarboxypeptidase |
|  | IV | Venom protease |  |  |
|  | VII | Serine protease snake | Serine protease 1-6 CUB<br>serine protease | Serine protease snake, CLIP serine<br>protease, CUB serine protease |
| *Esterase* | II | Venom acid<br>phosphatase Acph1-like | Acid phosphatase | Acid phosphatase 1 |
|  | V | Arginine Kinase |  |  |
|  | VI | Phospholipase A<sub>2</sub> | Phospholipase A<sub>2</sub>-1,<br>Phospholipase A<sub>2</sub>-2 | Phospholipase A<sub>2</sub>-1,<br>Phospholipase A<sub>2</sub>-2 |
| *Carbohydrate metabolism* | III | Hyaluronidase 1 | Hyaluronidase | Hyaluronidase |
| *Serine protease inhibitor* | VIII | Kunitz-type serine<br>protease inhibitor | Serpin 1-3 | Serpin 1-2 |

\* *B. sylvarum*, \*\* *B. humilis*

In *Thoracobombus* matchset 87 protein spots were manually matched in 2DE images of *B. humilis, B. pascuorum, B. sylvarum, B. ruderarius* and *B. zonatus* venom. Spot quantities were calculated and significant spots were detected statistically. Among these protein spots, 21 spots were present in each species. Common and differentially expressed protein spots are summarized (Table S2).

### 3.2. Mass-spectrometry of *Thoracobombus* Venom

From the 2DE gel images, we have spectrometrically characterized 47 proteins for *B. humilis*; 32 for *B. pascuorum*; 60 for *B. ruderarius*; 39 for *B. sylvarum* and 35 for *B. zonatus* (Fig. S1 and Table S3-7). Putative toxins are categorized according to their function and subcellular location (Van Vaerenbergh et al., 2015) (Figure 1). Toxins previously identified in *B. terrestris* and *A. mellifera* venoms are used as references (Table 2) (Van Vaerenbergh, Debyser, Devreese, & de Graaf, 2014; Van Vaerenbergh et al., 2015). Major putative toxins; phospholipase A2, venom protease, venom acid phosphatase and hyaluronidase were detected in venom of all five analyzed *Thoracobombus* species. Arginine kinase, a known component of spider and parasitoid wasp venom, was also identified in all studied species. Detection of serine protease snake (*B. ruderarius* and *B. sylvarum*), peptidases (*B. humilis* and *B. sylvarum*), and kunitz-type serine protease inhibitor (*B. humilis* and *B. sylvarum*), were limited to certain species.

### 3.3. Species Specific Expression of Putative Venom Toxins

Next, we compared the intensities of 2DE gel spots corresponding to putative toxins in each sample (Fig. 2). According to these spot intensities, several toxins display species specific expression patterns (Fig. 3). The expression of phospholipase A2, is the lowest in *B. ruderarius* venom among studied species. *B. zonatus* venom displays the highest venom acid phosphatase expression. *B. pascuorum and B. ruderarius* are characterized by a relatively low venom acid phosphatase expression. *B. humilis and B. pascuorum* display the most and the least abundant arginine kinase profile, respectively. *B. humilis* displays the highest venom protease expression. Hyaluronidase 1 expression is more than two folds higher in *B. ruderarius* venom compared to the other species.

**Fig. 2.**
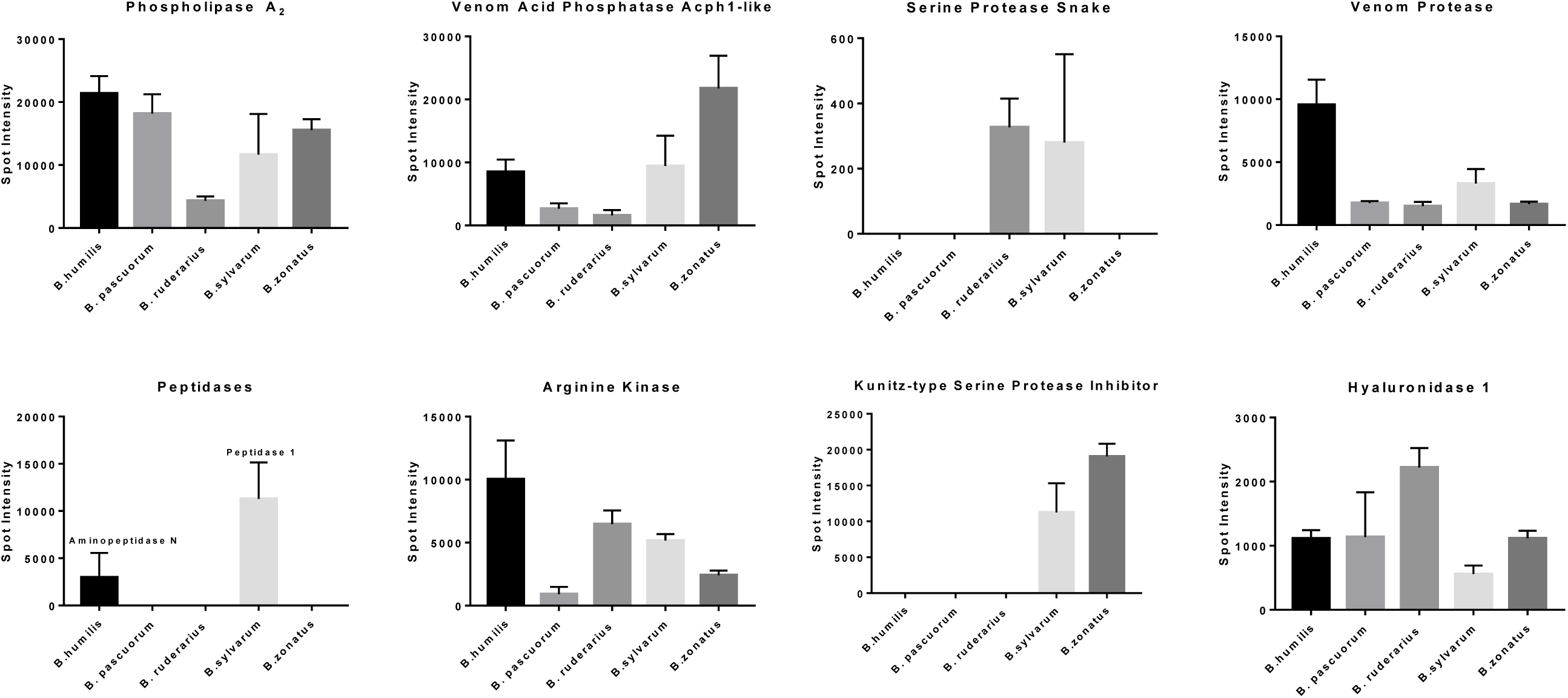
Relative 2DE gel spot intensities of putative toxins in *Thoracobombus* venom. Intensities, as calculated by PDQuest Software, are presented as mean ± standard deviation (3 replicates).

## 4. Discussion

Identification of *Thoracobombus* subgenus represents a challenge as its members are morphologically similar. In our previous study, we had applied landmark based geometric morphometrics to distinguish its members from each other more accurately. In this current study, we provide the primary proteomic characterization of the venom profile of five bumble bee species belonging to this subgenus. Several putative venom toxins are detected in *Thoracobombus* which were in accordance with *B. terrestris* and *A. mellifera* venom. Furthermore, we outline the species specific expression patterns of putative toxins.

In Hymenoptera, the defensive behavior of social groups is similar. Venom is used for protecting the colonies rather than for capturing prey. Therefore, the venom composition of these groups are expected to have common overall pattern. It is recently illustrated that the venom profile of *B. terrestris* and *A. mellifera* largely overlap, while also retaining certain species specific differences.

Accordingly, in *Thoracobombus* venom we have identified putative venom toxins collectively used by several Hymenoptera species to debilitate predators such as; phospholipase A2, arginine kinase, acid phosphatase (venom acid phosphatase Acph-1), serine protease (serine protease snake), peptidases (peptidase 1 and aminopeptidase N), hyaluronidase (hyaluronidase 1) and serine protease inhibitors (kunitz-type serine protease inhibitor).

While the *Thoracobombus* venom composition may be similar to other Hymenoptera species in terms of major toxins identified, it is important to note that the venom may be tailored for the individual species’ needs with regards to abundancies of protein families present. Accordingly, a change in the protein family profile of snake venom has been described upon nutritional shift.

Here, we have compared the 2DE gel images obtained from different *Thoracobombus* species and highlighted the species specific expression differences of venom proteins. In particular, our results indicate that expression levels of putative toxins may be used in differentiating between species. Within the *Thoracobombus* subgenus, *B. humilis* is morphologically very similar to *B. pascuorum; while B. ruderarius* and *B. sylvarum* resemble each other in terms of morphology and coat color. On the other hand, *B. zonatus* with its characteristic yellow coat color is easily distinguished from others. We have collected *B. ruderarius* and *B. sylvarum* at the highest altitudes (>2000m) and *B. zonatus* at the lowest (∼ 1000m). Interestingly, we have found that morphologically similar species differentiate in their expression of putative venom toxins, even in a similar habitat.

In future studies, different populations of same species can be used to assess the direct effect of altitude, vegetation, and prey population on the venom composition of bees. In this study we utilized 2DE gel images and MALDI-TOF mass spectrometry for assessing the expression levels of putative toxins, which may have its disadvantages. For example, we could not detect bombolitin, unique bumble bee peptides, in our samples. As bombolitins are very small peptides (10 kilo Daltons), we may have failed to detect these peptides with our equipment. More sensitive quantitative mass-spectrometry approaches may be necessary to confirm the correlation between microenvironment and the venom profile of bumble bees.

Profiling bumble bee venom can have pharmaceutical benefits. Venom peptides can be orally active and transported through the blood-brain barrier and through mammalian cell membranes. Commercial venom-derived drugs are available for the treatment of major diseases such as diabetes, stroke, and hypertension (King, 2011). Bumble bee venom may have the potential to serve as an alternative source of these drugs which are mostly snake venom-derived. Understanding the species specific expression patterns of toxins can also benefit the specific-compound oriented pharmacological analyses.

In conclusion, we provide the primary proteomic characterization of a diverse bumble bee subgenus, and illustrate species specific differences in toxin expression.

## Acknowledgments

This study was part of the PhD thesis of Nezahat Pinar Barkan. The experiments of this study were performed completely in Ankara University Biotechnology Institute. We are grateful to Selen Peker, Naşit Iğci, Beycan Ayhan and Hatice Yildizhan for their great help during laboratory work. We would especially like to thank Seçil Karahisar Turan for her great support and guidance throughout the study.

